# Generic versus personalized foot-ground contact models for predictive simulations of walking: Is personalization worth the effort?

**DOI:** 10.64898/2026.04.16.719049

**Authors:** Spencer T. Williams, Geng Li, Benjamin J. Fregly

## Abstract

**Purpose:** Quantification of walking function, including joint motions, ground reactions, and joint loads, outside the lab is a growing research area. Because only joint motions can currently be measured outside the lab, researchers are utilizing tracking optimizations of walking to estimate associated ground reactions and inverse dynamic joint loads. However, foot-ground contact models used in such optimizations have been generic rather than personalized, which may limit the accuracy of estimated ground reactions and joint loads. This study compares the predictive capabilities of generic versus personalized foot-ground contact models.

**Methods:** Generic and personalized foot-ground contact models were evaluated in calibration and tracking optimizations performed using experimental walking data collected from three subjects in varying states of health. Foot-only calibration optimizations evaluated how well both models could reproduce experimental ground reaction and foot motion data while tracking both types of data simultaneously, while whole-body tracking optimizations evaluated how well both models could reproduce experimental ground reactions, joint motion, and joint load data while tracking only experimental joint motion data and achieving dynamic consistency.

**Results:** For all three subjects and both types of optimizations, personalized foot-ground contact models reproduced experimental ground reaction, joint motion, and joint load data more accurately than generic foot-ground contact models.

**Conclusion:** Personalized foot-ground contact models can improve the accuracy with which ground reactions and joint loads can be estimated via tracking optimizations of walking using only experimental motion data as inputs. Personalized models require little time and effort to calibrate using freely available software tools and should improve the accuracy of predictive simulations of walking as well.

## Introduction

Researchers have used experimental data collected in a gait laboratory to study a wide range of movement impairments, including those caused by stroke [1], pelvic sarcoma [2], knee osteoarthritis [3], cerebral palsy [4], Parkinson’s disease [5], lower limb amputation [6], and incomplete spinal cord injury [7]. Commonly collected data include motion capture and ground reaction, which have enabled the calculation of joint motions and joint loads derived from those quantities. However, there is growing interest in taking experimental data collection outside the laboratory. The experimental hardware typically used in a gait laboratory is often expensive [3], people perform activities of daily living differently in a laboratory setting compared to their actual lives [8], [9], and patients could receive objective feedback on therapies ideally performed outside a clinical setting [10], [11].

Recent research advances are making it possible to take human movement measurement outside the laboratory, though significant limitations remain. Significant progress in hardware and software development (e.g., IMU systems [12] and markerless video motion capture [3], [13]) has made joint angle data available “in the wild”. Unfortunately, ground reaction data are more difficult to collect outside the laboratory. While instrumented shoe insoles exist [14], [15], [16], [17], [18], they do not capture all components of ground reaction forces and moments, plus most remain confined to a laboratory setting. Instrumented shoes have also been developed which can measure ground reaction forces and center of pressure, but the design influences subjects’ walking motions [19]. Without reliable ground reaction data, joint loads cannot be calculated via inverse dynamics.

Researchers are seeking to overcome these limitations by using computational methods to predict ground reactions and joint loads from motion data. Artificial neural networks can predict joint moments directly from more easily measured quantities like motion and electromyography (EMG) data [20]. Other approaches use measured motions and foot-ground contact (FGC) models within full-body predictive simulations of walking. These simulations typically use Hertzian contact models with Hunt-Crossley damping [21] to estimate ground reaction forces and moments between the foot and the ground. While the contact force equations in these FGC models are well-established, the parameter values used in these equations are almost always generic [22] and not validated to reproduce experimentally measured ground reactions from specific subjects. Furthermore, these simulation studies almost never report ground reaction moments, which are critical for calculating joint moments. Consequently, joint loads of clinical interest estimated using these models can be unrealistic for some subjects [3].

A likely explanation for unrealistic joint load results is the lack of foot-ground contact models calibrated to subject data. Historically, calibrating such models to actual data has been challenging because of the large number of parameter values that must be tuned (e.g., stiffness, damping, and friction) [23]. However, the Neuromusculoskeletal Modeling (NMSM) Pipeline [24] now makes it possible to calibrate personalized foot-ground contact models quickly and easily using experimental ground reaction and foot motion data. This automated calibration process removes the need for manual tuning of contact parameters [23]. When used in predictive simulations of walking, FGC models calibrated using this software have closely reproduced experimental measurements of all six ground reaction components [1], [24]. Personalized FGC models have never been more accessible, but how they compare to widely-used generic FGC models [22] for predicting ground reactions and joint moments from kinematic data has not yet been evaluated.

This study compared the predictive capabilities of generic versus personalized foot-ground contact models using two approaches. The first approach performed calibration optimizations that tracked experimentally measured ground reaction and foot kinematic data to test the models’ ability to reproduce both types of data simultaneously. The second approach performed tracking optimizations that estimated ground reaction forces and moments and lower-extremity inverse dynamics joint moments during walking from kinematic data alone while solving for dynamic consistency. For a robust comparison, the study was performed using experimental video motion and ground reaction walking data collected from three subjects in different states of health. Personalized FGC models have potential research applications inside and outside the laboratory and may help clinicians design better treatments for various movement impairments and assess treatment outcomes.

## Materials and Methods

### Experimental Data Collection

This modeling study used previously collected walking data obtained from three subjects. Subject 1 was a healthy male (age 47 years, height 1.7 m, mass 66.7 kg) [25]. Subject 2 was a male stroke survivor (age 79 years, LE Fugl-Meyer Motor Assessment 32/34 pts, right-sided hemiparesis, height 1.7 m, mass 80.5 kg) [1]. Subject 3 was a male with a right-side pelvic sarcoma who had not yet received surgery (age 46 years, height 1.73 m, mass 82.5 kg) [2]. All three subjects provided written informed consent, and data collection was approved by the institutional review boards of the University of Florida, Rice University, the University of Texas Health Science Center, and MD Anderson Cancer Center.

Although walking data were collected from the three subjects at different times and in different labs, the data collection protocols were nearly identical, supporting the use of these three data sets for the present study. All datasets included marker-based motion capture and ground reaction data. Motion capture data were collected using either a Vicon system (Vicon Corp., Oxford, UK) for Subjects 1 and 2 or a Qualisys system (Qualisys AB, Gothenburg, Sweden) for Subject 3, where marker placement utilized a modified Cleveland Clinic full-body marker set with markers added to the feet [26]. Ground reaction data were collected using a split-belt instrumented treadmill (Bertec Corp., Columbus, OH, USA) with belts tied for all three subjects walking at a self-selected speed (1.4 m/s for Subject 1, 0.5 m/s for Subject 2, and 1.0 m/s for Subject 3). One representative gait cycle with high periodicity starting with right heel strike was used for each subject. Subjects also performed a standing static pose to support skeletal model scaling.

### Skeletal Model Development

For each subject, a skeletal model with personalized lower-body joints was created starting from a generic full-body OpenSim [27] skeletal model [24] with 31 DOFs. The generic model possessed a 6 DOF ground-to-pelvis joint, two 3 DOF hip joints, two 1 DOF knee joints, two 1 DOF ankle joints, two 1 DOF subtalar joints, two 1 DOF toe joints linking the hindfoot and toes, a 3 DOF back joint, two 3 DOF shoulder joints, and two 1 DOF elbow joints. The model also included forearm pronation and knee adduction coordinates, though these coordinates were locked to values consistent with the standing static pose. The models were scaled based on the static pose motion capture data using the OpenSim Scale Model tool.

The Joint Model Personalization (JMP) Tool in the NMSM Pipeline was used to personalize select pelvis and lower body joint functional axes, body scale factors, and marker locations using each subject’s walking data, reducing inverse kinematics (IK) marker distance errors for gait motion data. Knee, ankle, and subtalar joint axis parameters, pelvis, femur, and tibia scaling, and femur and tibia marker positions were optimized. Joint angles were calculated by the OpenSim Inverse Kinematics tool for each subject’s gait motion using experimental marker trajectories and skeletal models updated by JMP. The resulting motion data were filtered with a fourth-order zero phase lag Butterworth filter with a cutoff frequency of 7/tf Hz, with tf equal to the duration of the gait cycle for each subject [28].

### Calibration Optimizations

Calibration optimizations were performed for both types of contact models using the Ground Contact Model Personalization (GCP) Tool within the NMSM Pipeline. These optimizations were used to evaluate how well both models could reproduce experimental ground reaction and foot motion data while tracking both types of input data. Prior to use in calibration optimizations, ground reaction data were filtered with a second-order zero phase lag Butterworth filter with a cutoff frequency of 7/tf Hz. For computational speed, foot models were automatically extracted from full-body models and hindfoot kinematics were redefined with a 6 DOF ground-to-hindfoot joint. This step avoided the need to account for lower limb geometry above the foot when making slight adjustments to foot kinematics during optimizations. For each type of contact model, the GCP cost function penalized errors in tracking ground reaction forces and moments as well as foot kinematics.

For personalized FGC models, the GCP tool evaluated how well measured ground reaction and foot kinematic quantities could be tracked simultaneously while automatically creating personalized contact geometry. A grid of viscoelastic contact elements with linear springs and nonlinear damping [21] was placed under each hindfoot and toes body using 37 locations within the bounds of each foot. The GCP tool determined the values of physical contact model parameters during these runs, including stiffness coefficients unique to each contact element and damping, dynamic friction, and resting spring length values common to all contact elements. As the personalized models had varying stiffness coefficients, the cost function additionally penalized Gaussian-weighted deviations in stiffness coefficients to ensure that nearby contact elements had similar stiffnesses without physically unrealistic discontinuities. Although stiffness coefficients were element-specific, they were shared between sides for each subject so that the contact models were symmetric. The optimization also adjusted the planar position of each force plate in the lab coordinate system to promote consistency between kinematic and ground reaction data [29], [30]. Ground reaction data with updated force plate positions were used to calculate joint moments with the OpenSim Inverse Dynamics (ID) tool. These joint moments were filtered with the same process as IK data, and they were used for evaluating the quality of predicted joint moments from simulations in future steps.

For generic FGC models, the GCP tool evaluated how well measured ground reaction and foot kinematic quantities could be tracked simultaneously while using generic contact geometry. Instead of automatically adding a grid of springs, models were manually created based on a generic contact model with six sparsely, nonuniformly placed spherical viscoelastic contact elements with nonlinear springs and nonlinear damping [21] commonly used in OpenSim [22]. Physical parameters of the generic model were not optimized; only minimized kinematic adjustments were allowed. These tests used the same ground reaction and kinematic tracking terms as the personalization process. The ground reaction data included the updated force plate positions from the personalization process for consistency.

### Tracking Optimizations

Tracking optimizations were performed for both types of contact models using the Tracking Optimization (TO) Tool within the NMSM Pipeline. For each subject, these optimizations were used to evaluate the ability of each type of contact model to predict ground reaction forces and moments while tracking only kinematic data and solving for a dynamically consistent walking motion. Coordinate positions, velocities, and accelerations were included as design variables, and torque controls were used for hip, knee, ankle, and subtalar coordinates. The cost function tracked only coordinate positions and velocities for all joints except the ground-to-pelvis joint, leaving ground reactions and joint moments to be predicted during the optimization. The constraint function prevented significant changes to initial coordinate positions and velocities. It also imposed dynamic consistency by keeping torque controls within 0.1 Nm of their respective ID joint loads during the optimization and by constraining the residual ID loads of the 6 DOF ground-to-pelvis joint to be less than 1 N or 0.1 Nm in magnitude.

## Results

### Calibration Optimizations

For all three subjects, GCP tracking errors were lowest when using personalized instead of generic FGC models (Table 1). Overall, kinematic and ground reaction quantities were roughly 2-3 times more accurate with personalized models. As a visual example, tracked quantities and stiffness coefficients for the right foot of Subject 1 are plotted in Figures 1 and 2.

**Table 1.** Root mean square error (RMSE) (mean ± standard deviation) in GCP tracked quantities on the right and left sides of all subjects for the calibration optimizations.

|  | Axis | Force (N) | Torque (Nm) | Translation (mm) | Rotation (°) |
| --- | --- | --- | --- | --- | --- |
| **Generic** | **X** | 17.4 ± 10.8 | 1.45 ± 0.65 | 7.77 ± 2.98 | 3.12 ± 0.72 |
| | **Y** | 8.82 ± 3.41 | 0.72 ± 0.39 | 12.3 ± 2.07 | 1.45 ± 1.04 |
| | **Z** | 5.67 ± 3.31 | 1.41 ± 0.39 | 3.00 ± 1.41 | 1.93 ± 0.49 |
| | **Toes** | - | | | 2.12 ± 0.93 |
| **Personalized** | **X** | 6.04 ± 2.79 | 0.74 ± 0.35 | 3.21 ± 1.05 | 1.30 ± 0.23 |
| | **Y** | 4.31 ± 1.67 | 0.39 ± 0.17 | 4.06 ± 0.78 | 0.85 ± 0.32 |
| | **Z** | 2.16 ± 0.73 | 0.83 ± 0.39 | 3.13 ± 1.11 | 1.24 ± 0.17 |
| | **Toes** | - | | | 0.85 ± 0.19 |

**Figure 1.**
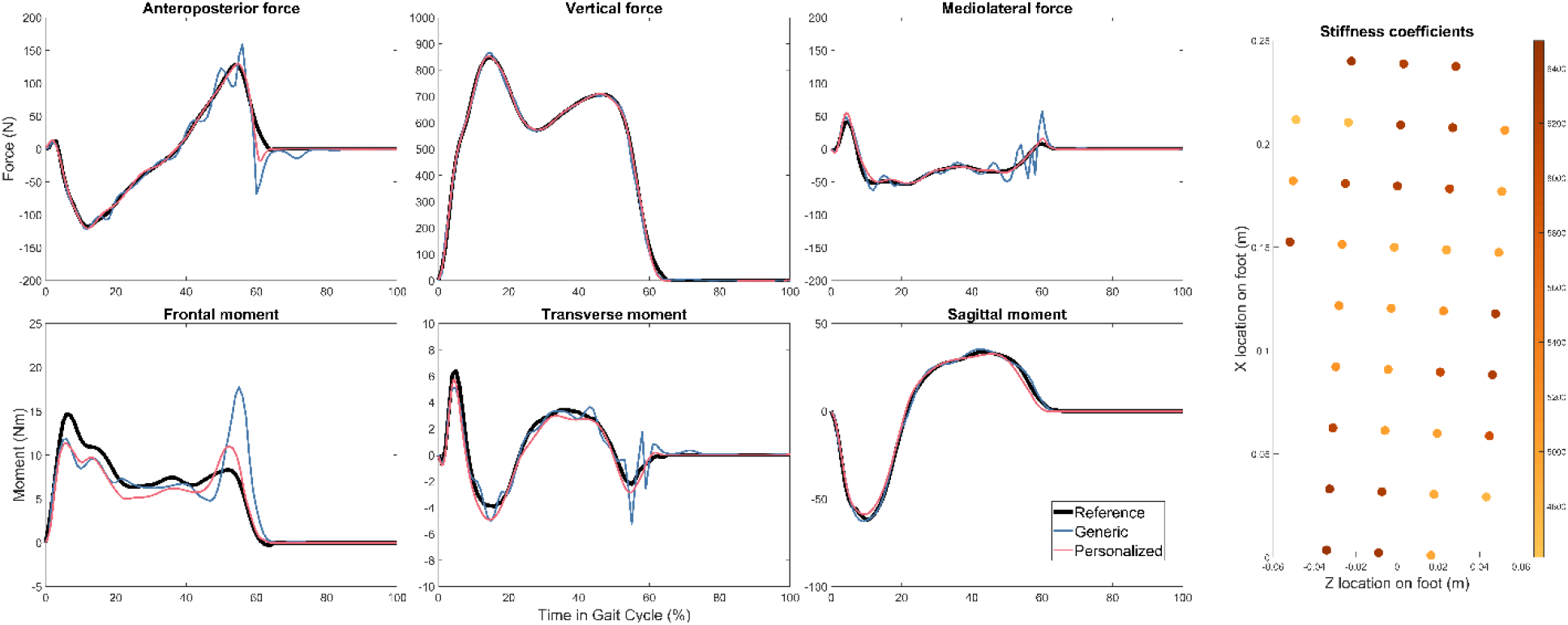
Right foot reference ground reactions and GCP calibration optimization results with generic and personalized FGC models for Subject 1. The distribution of stiffness coefficients on the personalized model is shown on the right.

**Figure 2.**
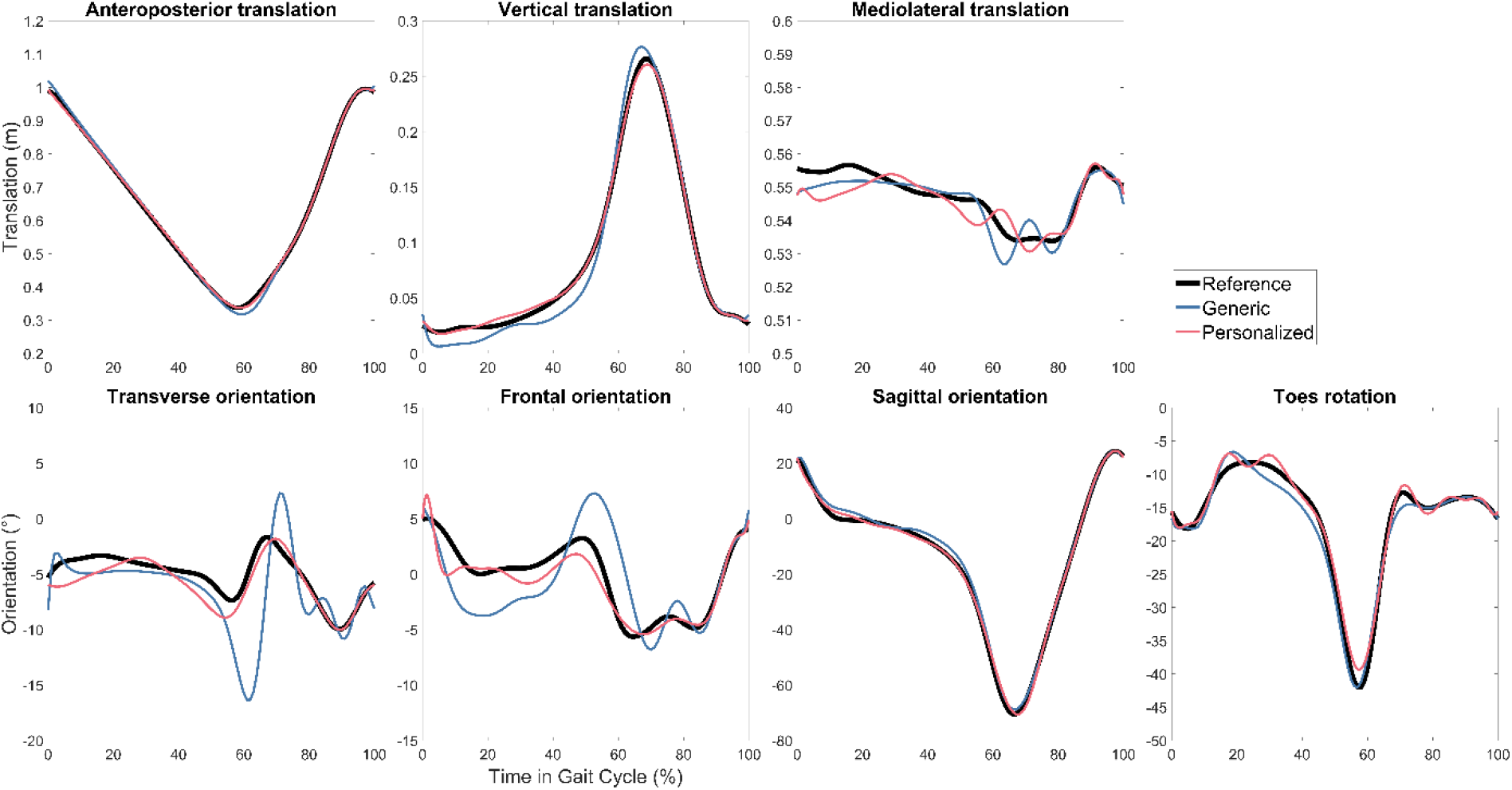
Right foot reference kinematics and GCP calibration optimization results with generic and personalized FGC models for Subject 1.

### Tracking Optimizations

For all three subjects, TO tracking and prediction errors were lowest when using personalized instead of generic FGC models (Table 2). Motion tracking was roughly 4-5 times more accurate with personalized models, while ground reaction predictions were roughly 3-4 times more accurate, and inverse dynamics load predictions were roughly 2-3 times more accurate. As a visual example, tracked motions, predicted ground reactions, and predicted joint loads for Subject 1 are shown in Figures 3, 4, and 5 respectively. Figure 6 shows the positions of skeletal geometry over the course of the gait cycle for Subject 1 for the nominal motion alongside motions predicted using each contact model.

**Table 2.**
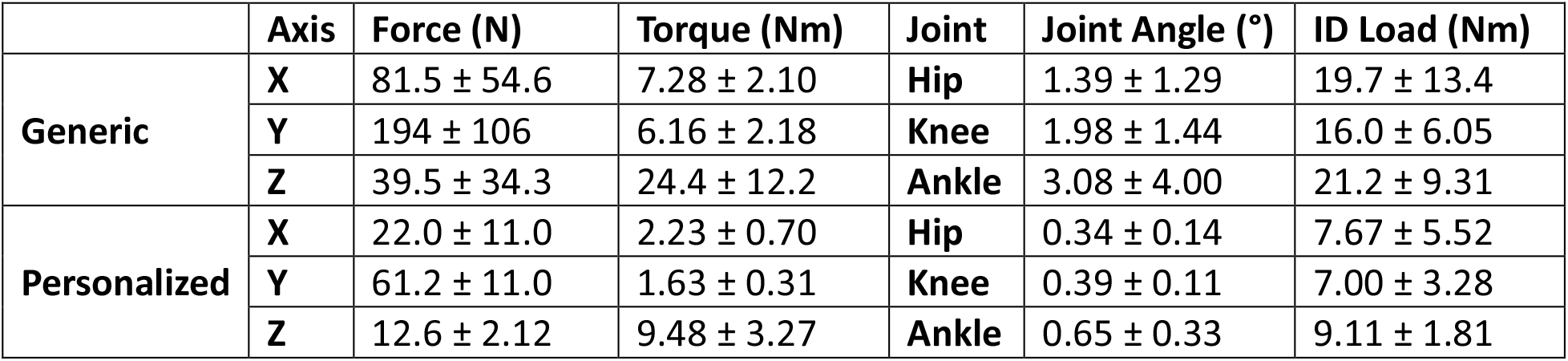
Root mean square error (RMSE) (mean ± standard deviation) in tracked and predicted quantities on the right and left sides of all subjects for the Tracking Optimizations. For joint-level quantities, the hip includes hip flexion, adduction, and rotation, the knee includes knee flexion (as well as knee adduction for ID loads), and the ankle includes ankle flexion.

**Figure 3.**
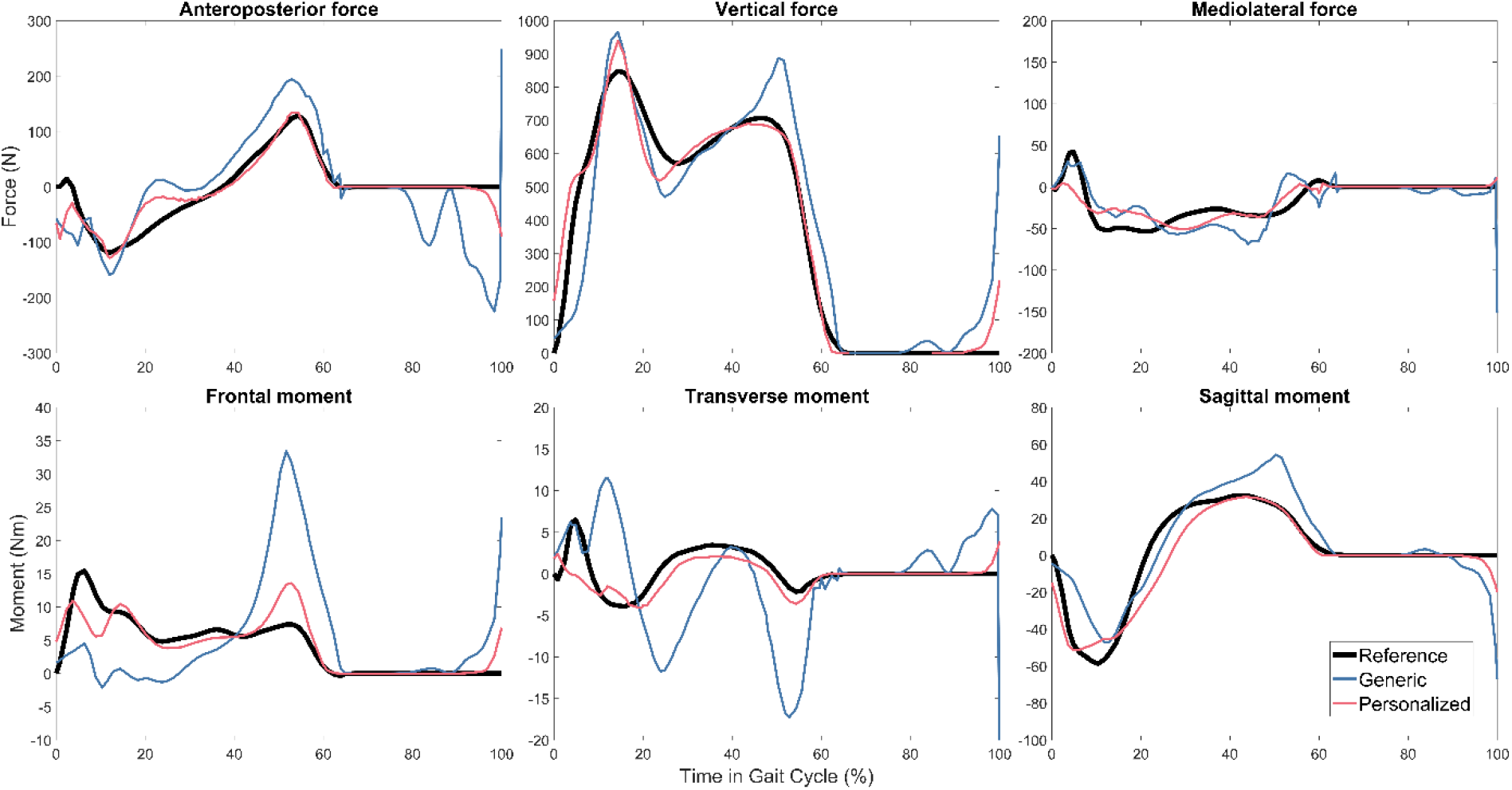
Right foot reference ground reactions and Tracking Optimization results with generic and personalized FGC models for Subject 1.

**Figure 4.**
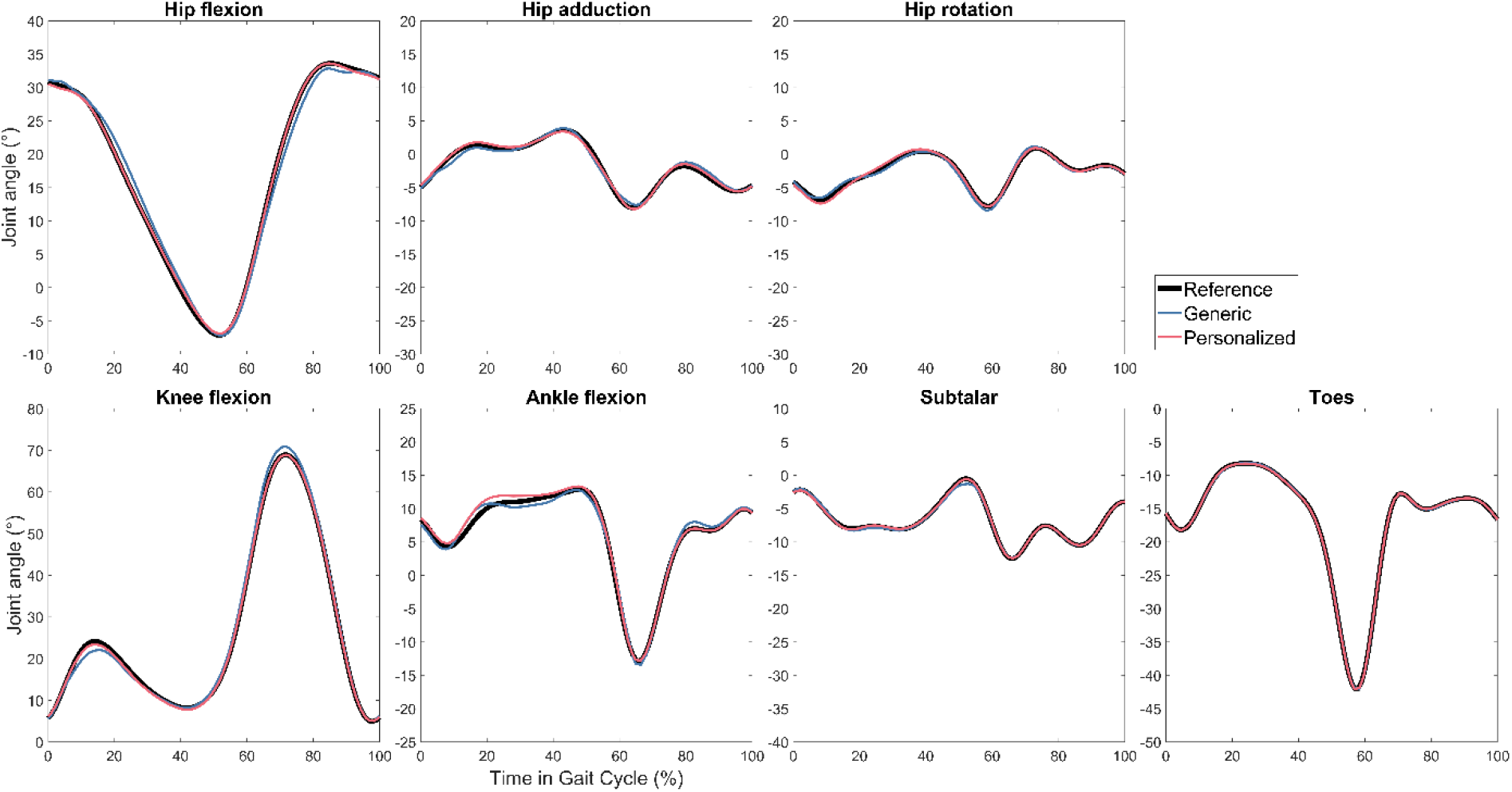
Right leg reference kinematics and Tracking Optimization results with generic and personalized FGC models for Subject 1.

**Figure 5.**
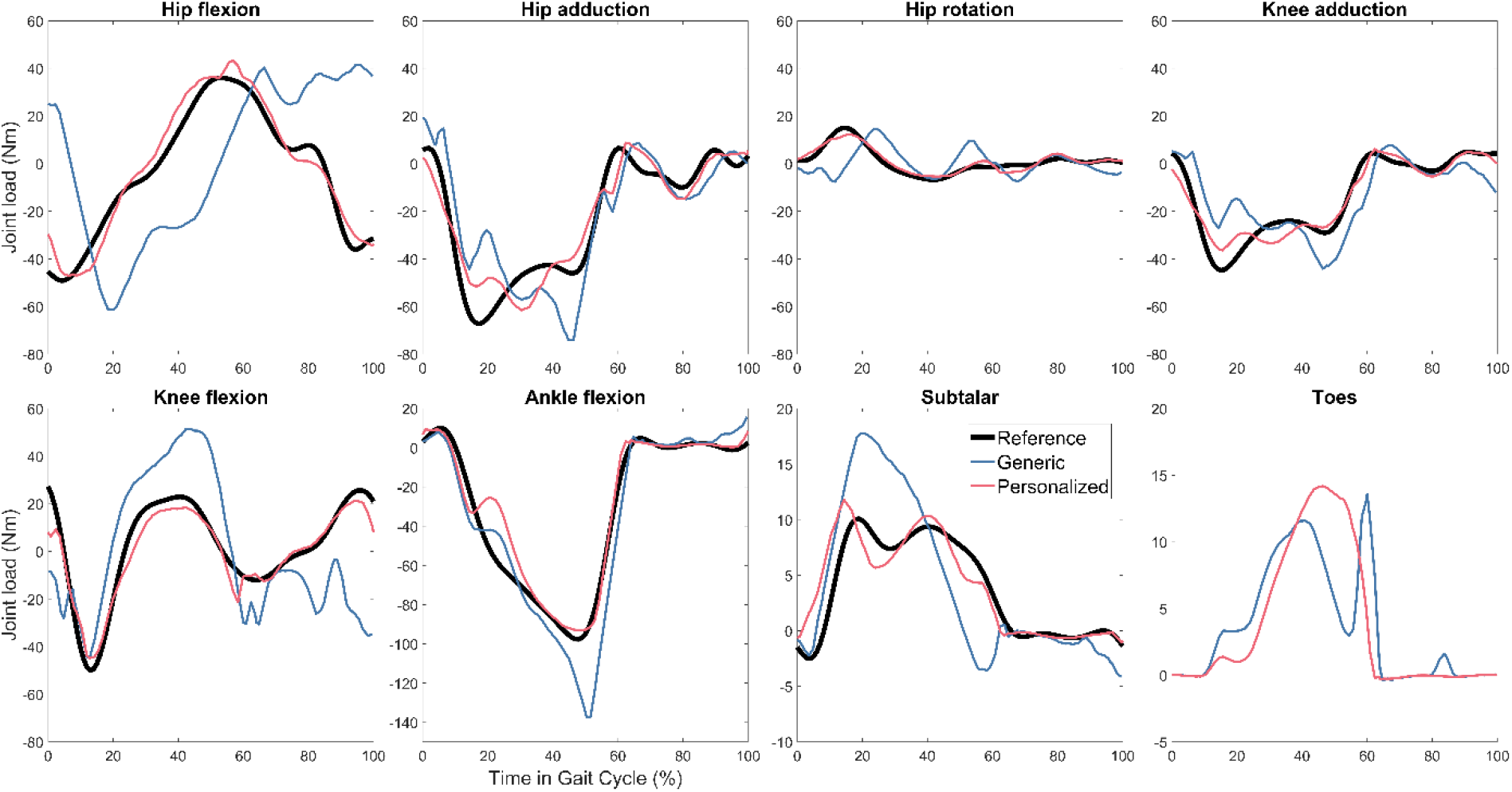
Right leg reference joint loads and Tracking Optimization results with generic and personalized FGC models for Subject 1. No reference load is plotted for the toes because the reference ID results did not have ground reactions distributed between the hindfoot and toes bodies, so there is no meaningful reference for this joint load.

**Figure 6.**
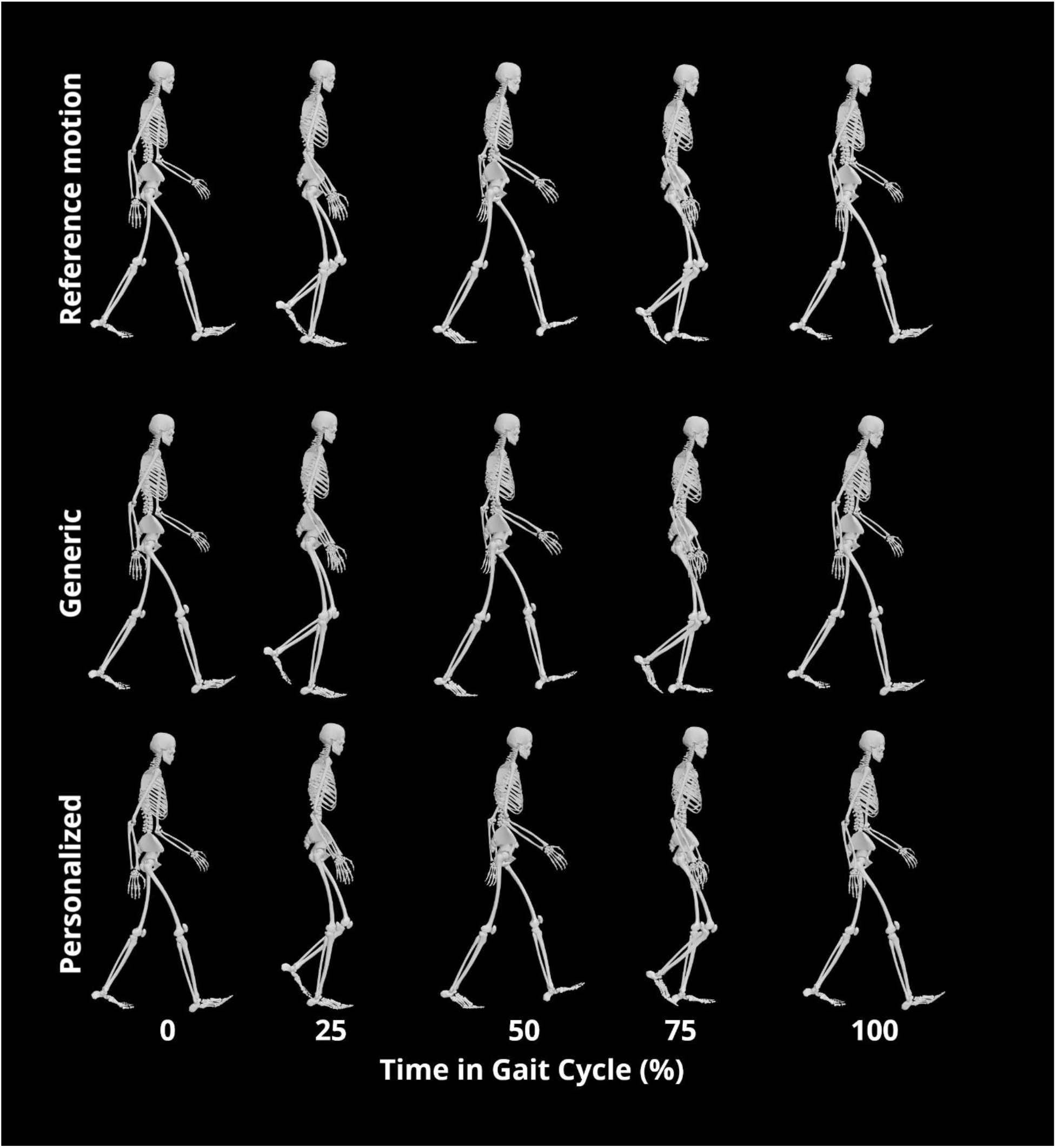
Animation strip of reference and resulting motions from Tracking Optimization for Subject 1. Note the differences in foot positions at 25% and 75% and in truck orientation at 100%.

## Discussion

This study compared the predictive abilities of generic and personalized FGC models for estimating ground reactions and joint loads and modeling dynamically consistent motions. The three subjects included provided a wide variety of test conditions, showing FGC model performance for a healthy subject, a stroke survivor, and a subject with a pelvic sarcoma. For all three subjects, the calibration and tracking optimization runs successfully converged for both generic and personalized FGC models. In calibration results, the personalized FGC models had lower RMS errors in all tracked quantities than did the generic models except in hindfoot Z translation, which differed on average by less than a millimeter. The improved performance of personalized models in calibration optimizations is an intuitive result since the physical parameter values were adjusted in the personalized models to reduce errors. However, it is more notable that the personalized models performed significantly better in tracking optimizations, greatly reducing errors in ground reaction and ID joint load predictions and also reducing errors in tracked joint angles when solving for dynamically consistent walking motions. These results suggest that although both generic and personalized FGC models can produce reasonable ground reactions from walking motions and can support dynamically consistent walking simulations, personalized FGC models have the potential to produce significantly more accurate predictive simulations for individual subjects.

While conducting this study, we made additional observations that may provide further insight into the use of FGC models for predictive simulations. First, FGC model personalization can reduce tracking optimization computation time. For all subjects, predictive TO simulations using personalized FGC models converged in fewer iterations than simulations using generic FGC models. For two of the subjects, tracking optimizations with personalized FGC models also converged in less time, saving more time than was required to calibrate their contact models. For the third subject, the tracking optimization with a personalized FGC model converged in a comparable amount of time. A speed increase in tracking optimizations would be especially beneficial for users intending to run multiple predictive simulations for the same subject. Second, a TO problem formulation more consistent with past work [3] produced unrealistic toe motions. This formulation included tracking joint accelerations and removing tracking from the toes coordinate to let it change freely. Acceleration tracking yielded slight reductions in ground reaction tracking errors, though joint angles were slightly less accurate. Allowing the toes coordinate to move freely also slightly (though less consistently) reduced ground reaction tracking errors overall. However, the toes moved in a highly unrealistic fashion, attempting to flex and extend by more than 20 degrees from their nominal motions.

The performance of our generic FGC model in tracking optimizations is consistent with past work, though it is important to note that the weighting of cost function terms is key to how errors in simulation results are distributed. For simulations predicting ground reactions, it is possible to have higher accuracy in ground reaction and joint load predictions at the cost of less accurate motion tracking [3], [31]. As seen in the present study, irregular shapes may appear when plotting ground reactions over time from results generated using generic FGC models [32], though these plots are not always available in published work. Similar errors are observed in other motions, such as sit-to-stand simulations [3], [33]. It would additionally be useful to compare the results of this study’s ground reaction moment predictions to past work. Use of personalized FGC models yielded dramatic improvements in predicted ground reaction moments, but they are often not reported, even for studies using three-dimensional models.

Though large differences in tracking and predictive errors between generic and personalized FGC models make the conclusions of this study clear, this work has limitations that are important to consider when applying these findings. First, the data used in this study represent a best-case scenario for both calibration and prediction. For all three subjects, the motion capture data came from a multi-camera marker-based system, which is currently the “gold standard” in the biomechanics field and the most accurate motion capture method without using ionizing radiation [34], [35]. For reasons such as budget and the ability to measure motion outside limited laboratory settings, researchers are becoming increasingly interested in other motion capture methods such as IMU and markerless video motion capture [3], [12], [13]. Both the GCP and TO tools are compatible with joint angle motion data derived from any motion capture method, so this study and its procedures can be applied to these motion capture methods as well. However, lower-quality motion capture data will likely lead to a lower-quality model calibration, so it is unknown how well an FGC model personalized using these data would perform. The TO tool would still need to resolve any dynamic inconsistency in the captured motion, so it is likely that using either a generic or a personalized FGC model with lower-quality motion data would produce higher predictive and tracking errors than those found in this study. Second, all simulations in this study were torque-driven, so muscles and their limitations on possible motions and control were not accounted for. The inclusion of muscles can be especially important for predicting new motions [1], so it will be valuable to confirm that the trends in the results of this study remain valid when using more complete musculoskeletal models. Third, FGC models were used in this study to predict dynamically consistent versions of the same motions used in calibration. It would be useful to evaluate the predictive capabilities of FGC models for new situations, such as walking at a faster speed or running.

In conclusion, we find that personalized FGC models more accurately predict dynamically consistent walking motions than do generic FGC models for subjects in a variety of states of health. The study demonstrates that the automatic personalization process for FGC models is robust, fast, and easy to perform for different subjects. With high reward for little cost, foot-ground contact model personalization appears to be worth the effort. Personalized FGC models may enable higher-quality predictive simulations in biomechanical walking studies of many movement impairments in an array of settings.

## Data Availability

Experimental data, results, and code used to generate the results of this study are available here on SimTK.org.

## Contributions

Spencer T. Williams performed all neuromusculoskeletal modeling work after Joint Model Personalization, formulated and ran all other NMSM Pipeline tool runs, and participated in writing the manuscript draft. Geng Li performed initial neuromusculoskeletal modeling, including model scaling and Joint Model Personalization, and processed all data used in this study. Benjamin J. Fregly supervised this project, assisted with optimal control problem formulations, evaluated optimal control results, and helped with revising the manuscript draft.

